# A Graphic User Interface (GUI) to build a cost-effective customizable 3D printed Prosthetic Hand

**DOI:** 10.1101/2020.03.18.997486

**Authors:** J. Lázaro-Guevara, R. Gondokaryono, L. González, K. Garrido, N. Sujumnong, A. Wee, J. Miscione

**Affiliations:** Worcester Polytechnic Institute, Worcester, MA, USA; University of San Carlos of Guatemala, Guatemala; University of Utah, Salt Lake City, UT, USA

**Keywords:** 3D printing, CAD design, customizable prosthetic hand

## Abstract

This project aims to create a tool that allows medical staff to use an intuitive graphical user interface (GUI) based application to generate STL models of a customizable prosthetic hand, that can be 3d printed to fit a specific patient’s hand size. Since the whole process of adjust and adapt the prosthetics devices could consume most of the resources of small medical attention centers. And the necessity to adapt the prosthetic devices is highly relevant when these are intended to be used by the pediatric population. This software creates a customizable parametric 3d model for a trans-radial prosthetic hand and all its necessary components for 3d print and assemble it. The software is intended to be operated by non-trained staff, reducing the costs of remodeling or adapting the original model to fit the necessity of a patient, allowing to produce personalized prosthetic devices in a cost-effective manner with an effortless customization approach. This will allow that medical practitioners with a lack of technical background to get involved in spreading 3D-prosthetics. Also, using open-source parametric 3D-models could lead to existing 3D-prosthetic projects that will adopt this method of customization, allowing the expansion of 3D-printed prosthetics at developing-countries reaching all needing patients. Ultimately, this tool will allow the medical staff to focus on adjusting or replacing the prosthetic devices more often than previously, due to be considered too expensive..

## BACKGROUND

Loss of arm function due to amputation from combat, birth defects, or accidents negatively affects amputees in terms of independence. Additionally, many amputees cannot afford the prostheses that are available due to the high cost. This is especially true for young children, as they grow quickly and require more frequent replacement^1-3^.

The focus of this project is to construct a trans-radial prosthesis that is affordable (sub $100 range). A trans-radial prosthesis is an artificial limb that is a replacement for below-the-elbow amputation. Generally, there are two classes of functional (non-cosmetic) prostheses in this category: a hook-and-loop cable driven arm, and an electrically driven arm^2, 4, 5^. A patient may also choose to have a cosmetic arm (which has no actuation functionality)^2, 6^. Table I illustrates the price range that a patient (or insurance company) would have to cover in order to have an artificial limb ^7, 8^.

**TABLE I.** TYPICAL PROSTHETIC ARM COSTS

| Type of Arm Prosthesis | Price Range (\$) |
| --- | --- |
| Cosmetic arm | \$1,000 - \$5,000 |
| Hook-and loop arm | \$5,000 - \$10,000 |
| Myoelectric arm | \$10,000 - \$100,000 |

Despite many mechanical advances, prosthetic arms, although lighter and stronger today than ever before, are not intuitive to control. Certain groups have been working towards better control feedback methods, such as Touch Bionics or DEKA. Touch Bionics uses a traditional two-input myoelectric (muscle signal) that is detected by electrodes sitting on the surface of the skin to open and close their hands grip. DEKA created a modular arm usable by all arm amputees and for full arm amputees. The user undergoes a surgery to move the nerves from the upper spinal cord across the shoulder, and over to the pectoral muscles, which when they contract to send electrical impulses through skin-mounted electrodes on the chest ^9-11^.

This same arm provides feedback through small vibrating motors that are mounted to the user’s skin. The vibration signal changes with grip strength. As the user’s grip tightens, the frequency of the vibration increases^10^. However, neither of these solutions seems possible to be applied to current prosthetic devices in an affordable way. Besides detecting electrical impulses, another method that is used to control prosthetic arms is via pressure sensors located in the shoe ^11^. Although not exactly intuitive to use, it did seem like a more reliable solution than EMG sensors. Another patent filed by DEKA shows that they developed a system that could command a prosthetic arm via sensors that picked up the user’s movement ^3, 12^.

With the onset of technological improvements such as 3D printing and smaller sensor packages, it is now possible for individuals to design a personalized prosthetic arm in a cost-effective way ^7, 13^. Some of these are mechanically actuated (driven off wrist-joint movement), such as the open-sourced designs by Enabling the Future. They generally tend to be in the $500-$2000 price range if they are electric and $50-$1000 for purely mechanical arms^1, 2, 8, 14^. For this to be a viable option for amputees in developing countries, the hand must cost less than $100 total ^15^. Not taking in consideration, the cost of skilled labor in producing such medical devices as prosthetics that may be adversely affect the production cost on developing economies^6, 16^.

## I. THE PROJECT

### A. Overall Goal

The purpose of this project is to create a software tool that generates STL files for a prosthetic hand based on input of patient medical data without the direct use of any CAD software. In addition, the tool will produce a BoM required for assembly. The system must be tested to ensure that it creates designs that can withstand grasp force minimums without unnecessarily straining the actuation motors. In order to achieve this, the specific STL files will be loaded into a simulation program to determine if their geometry can complete expected tasks.

To accomplish this overall goal the next steps are necessary to succeed on this project.

1. The design and application of all dependencies created using completely open source software (Open SCAD).
2. All automated generated prosthetic hands (STLs) will be assessed to quality-control analysis to ensure every prosthetic generated file meets desired benchmarks.
3. A test generated STL file will be physically printed and subjected to case scenarios ensuring feasibility of the project.

### B. Design main considerations

#### Mechanical Optimality

– Reducing the amount of 3D printed material required to achieve an acceptable safety factor under operating conditions lowers the device cost and decreases the time required to print components.

#### Actuation Optimality

– Electromechanical designs have the drawback of requiring the use of batteries. The more the user can use the device without needing to recharge makes the prosthetic more successful.

#### Implementing Cost

– Using as much open source software as possible allows medical staff to focus spending on quality materials rather than necessary software.

## II. METHODS

This section includes descriptions about software, CAD, mechanical, and 3D implementation.

### A. Obtaining the Input parameters to the GUI

The input parameters to the GUI will be introduced by a previously trained physician who will provide information on the specification of the prosthesis for individual patients, these measures will be previously obtained by the medical staff. The necessary input to the GUI will be formatted in drop-down boxes (for questions) or edit fields (for measurements) on the GUI.

The flow from collecting data to fitting a 3D printed prosthetic hand is described in Figure 1.

**Fig. 1.**
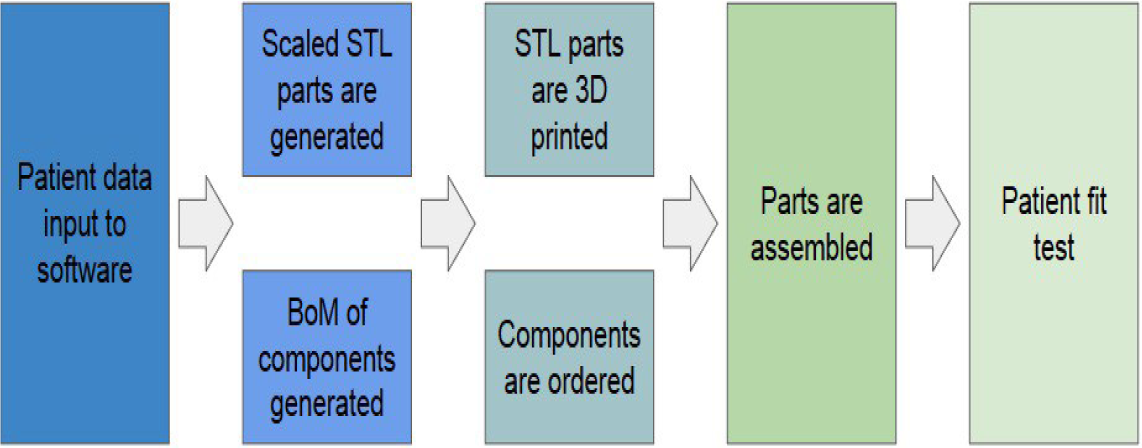
Flowchart showing the operation sequence of the system

### B. GUI interface

The software is written in Python and uses **Tkinter** for GUI development. Python is capable of running .bat scripts which can be used to automate the STL generation as described in the CAD Parametric Design section.

The GUI provides medical staff with several parameters to enter based on the patient as seen below in Figure 2. Currently, all parameters are input individually.

**Fig. 2.**
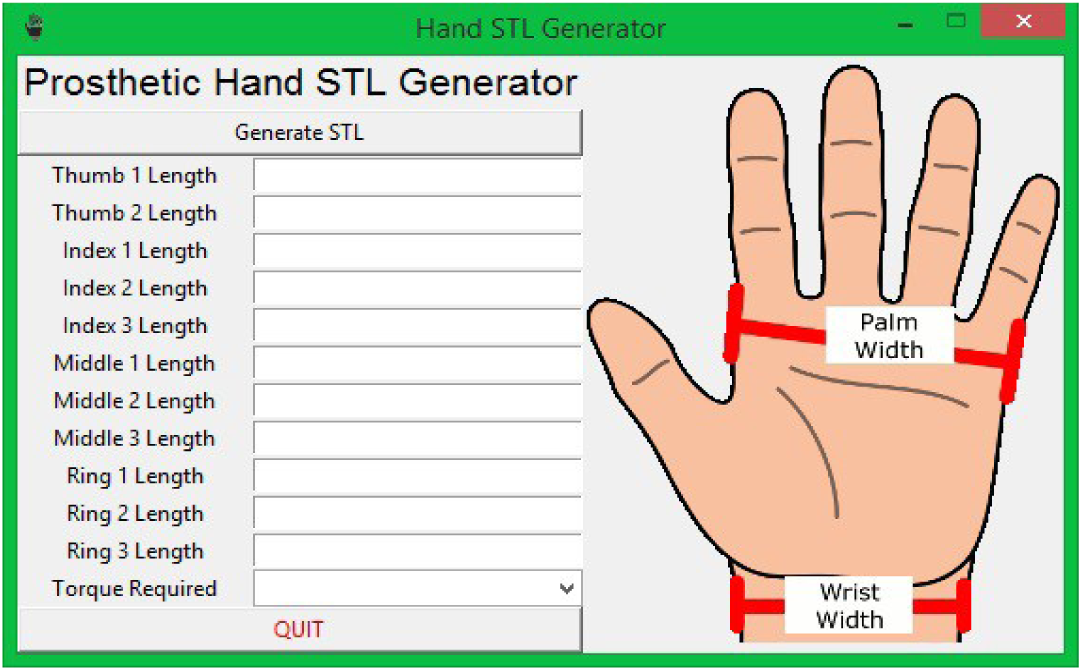
The preliminary GUI allows the user to enter in specific measurements about the patient’s hand.

STL creation is based upon modifying OpenSCAD files, since OpenSCAD is a text based language, modifying the files modifying the parameters of the CAD construct is an easy task to accomplish this we developed a text Edit function that sets the first line of each .scad file to contain the variables needed to correctly scale the model.

Once the .scad files are created, they are converted into STL files using a batch file (obtained from http://jsfiddle.net/Riham/yzvGD/embedded/result/). This batch file is called from the Python application and runs without starting the OpenSCAD interface. Unfortunately, the batch file requires the OpenSCAD directory to be specified.

Aside from creating the STL files, the application also creates a bill of materials that can be customized to each user specification. Currently, the BoM only produces a CSV file with motor selections based upon the “Torque Required” selection box of the GUI. The GitHub repository for the application can be found at (https://github.com/megahitokiri/A-Graphic-User-Interface-GUI-to-build-a-3D-printed-personalized-Prosthetic-Hand)

### C. CAD parametric Design (OpenSCAD)

In order to reduce costs and keep the STL files opensource the code will be designed to utilize OpenSCAD for parametric modeling. OpenSCAD’s file structure is essentially high-level language text file.

This can be created entirely by Python and exported to the scad format to later be converted to STL. The image generated by openSCAD can be seen in Fig 3.

**Fig. 3.**
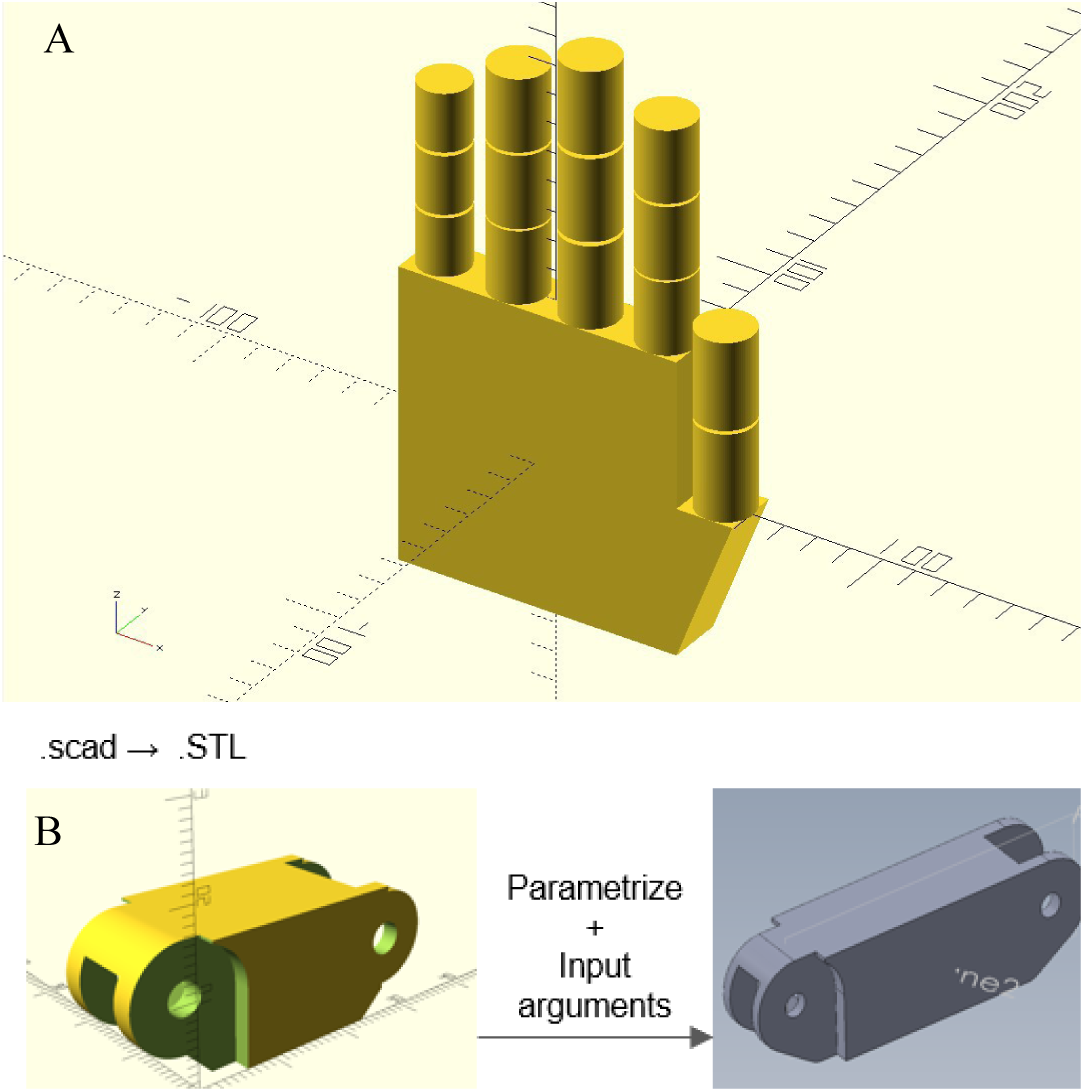
A. The concept parametric design in openSCAD B. From OpenSCAD rendering to STL to 3D visualization.

Once the core structure of the hand is generated, the SCAD model will be exported in STL format for appearance reconstruction to make the hand looks aesthetic.

### D. CAD parametric Design (OpenSCAD)

We use a multi-pulley system on the palm to actuate close five fingers with one motor. Standard 5/32” Dia. Dowel Pins and 0.188 OD Diameter Torsion springs have been chosen for all hand joints.

For different hand width, the length of the Dowel pins will change. Also, there are two possible 0.188” torsion springs (one with 3 coils and another with 7 coils) that will be chosen based on force of grasp and/or width of finger. 0.093” Dia. Dowel Pins have been chosen for the guides of the cable through the palm and the fingers Figure 4A.

**Fig. 4.**
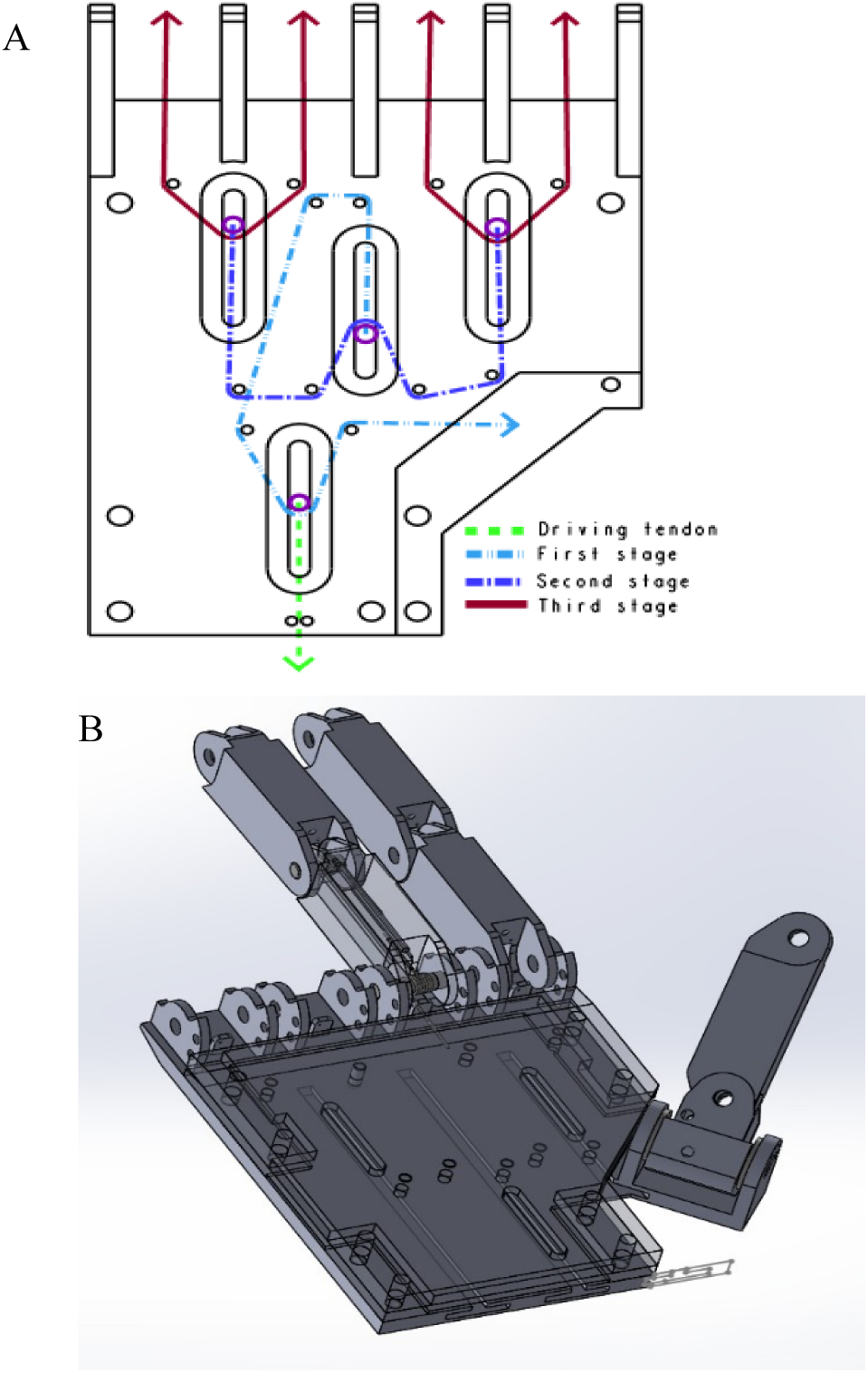
A. Pulley System B. Partial hand design in SolidWorks

The Palm is comprised of three parts. The bottom layer is for the slider of the multi-pulleys. The slider itself will be a 3D printed T-joint with pin. The second layer is for the guide of the slider to stop at a certain point and keep perpendicularly straight.

The third layer palm cover layer is to keep the pulley dowel pins in place, cover the cables, and as an aesthetic part of the palm of the hand. The thumb joint angle can be manually adjusted with a 4-40 screw to accommodate different grasps. It is though still actuated with cable from the same pulling force.

The Mechanical Design Fig. 4B performed in SolidWorks is later extrapolated onto OpenSCAD, this adds aesthetics of the prosthetic hand to look more like a human hand. Aesthetics will be added on the finger joints, the back of the hand, and the palm of the hand.

### E. Anthropometric Measurmetns Adjustments

Anthropometry is an important part of every mechanism that is intended to be used by a patient, ranging from the ergonomic analysis to the functional analysis. A normal value varies between different populations but is also impacted by inter-subject differences^17^. However, there are percentiles that are considered normal for the 90 percent of the population (5th to 95th percentile). Even though this prosthetic hand is designed to be personalized, it should not be outside this normal range as indicated in Table II. This range was calculated based on the database from the project AntroKids (sponsored by National Institute of Standards and Technology: http://ovrt.nist.gov/) that compiles measurements from young people ranging from age 3 to 20. We focused on the average values from ages from 7 to 20 (considering 20 as the terminal growing age for hand dimension) for making this prototype. These are some of the parameters the user assigns in the GUI.

**TABLE II.** NORMAL ANTROPOMETRIC MEASURES

| Measurement (in cm) | Mean | 5th % | 95th % |
| --- | --- | --- | --- |
| Hand Length (age 7) | 13.8 | 12.6 | 15.1 |
| Hand Length (age 20) | 18.4 | 16.2 | 20.2 |
| Hand Width (age 7) | 4.4 | 3.5 | 5.2 |
| Hand Width (age 20) | 9 | 8.6 | 9.5 |
| Middle Finger Length (age 7) | 12.2 | 11.1 | 13.2 |
| Middle Finger Length (age 20) | 15.8 | 12.9 | 18 |

## III. RESULTS

### A. 3D printing the GUI generated STL files

Using a Wanhao Duplicator 5S and PLA material at manufacturer recommended printing temperature ranges (190 to 210 C°) on heated bed and introducing at the GUI for 3D printing the 5th and 95th percentile standard measures for 20 years old patient (Table II). We proceed to 3D print the generated STL files from their OpenSCAD model Figure 3A.

From the previously parametrized values parsed from the GUI to the openSCAD algorithms the STL can be easily visualized and manipulated in any 3D printing STL software (e.g. Cura) figure 3B.

And the scaling representation from the model visualization step (3D rendering) to the printing steps varies in less than 0.5%, meaning that the accuracy of generating the models is inside the expected parameters as shown by figure 5A (95^th^ % ∼ 9.5 cm).

**Fig. 5.**
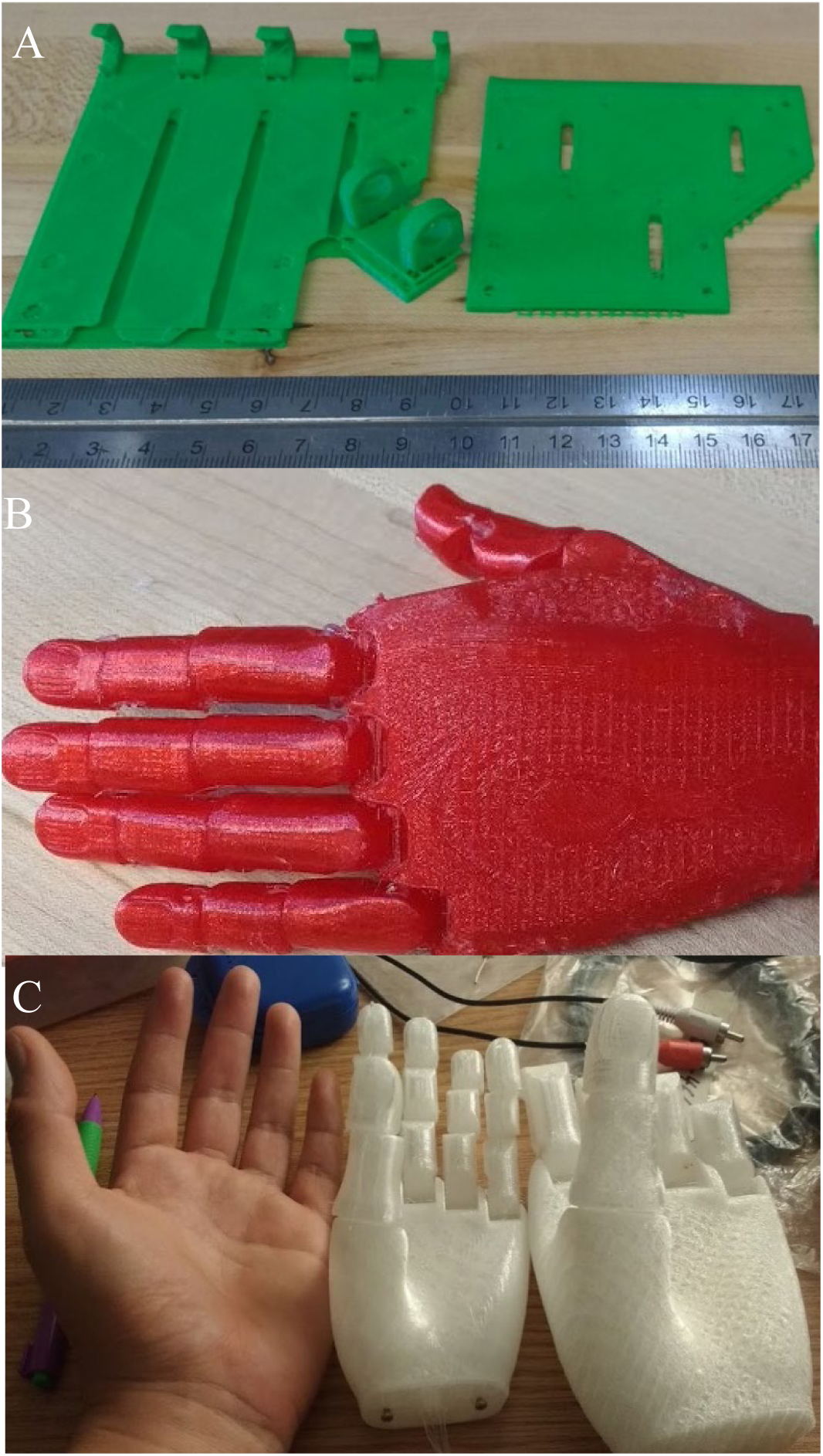
A. 3D Printed visualization of STL files 99% accuracy size. B3D Print using flexible filament size accuracy varies from stretching. B & C. Showing the customizability of parametric models for 3D printing.

This minimal variation could be due to the printing techniques and can be easily adjusted to the patient’s specifications due to the thermoforming properties of PLA that allows certain degree of bending when heated Figure 5B. When using a different printing material like Ninjaflex the aesthetics of the 3D model can be improved Figure 5A (PLA) and 5B (Flexible Filament). Also, when performing the 3D printing of the models by the GUI algorithm (5th to 95th %) there is a 1 to 1.5% scale variation against the expected outcome size (real 3D printed) against the inputted size at GUI and the generated parametric models Figure 5C.

This variation does not seem to correspond to an escalation error when analyzing the rendered models, instead it is likely that the discrepancies corresponds to an inconstant rate of deformation on flexible filaments on varied 3D printing conditions.

Because of the multiple complications of 3D printing using flexible filaments, these materials are not suitable for a non-expert user since the accuracy of the model could vary from the stretching of the material.

## IV. DISCUSSION

With the availability and cost reduction of 3D printers around the world. Tools that helps in making user friendly 3d models access to the population are necessary ^4^. And there is no better place to start making accessible models like the medical realm, where a simple customization feature can highly improve the quality of multiple patients.

Therefore, providing the ability to project and rotate virtual 3D object that later on can be translated to a 3D physical construct can help the physicians to get involved and start providing this kind of services to rural areas or not highly specialized hospital were many patients can benefit from ^13, 15^.

Also, making these tools Free and open-source will make them evolve quicker and with and easy availability to all kind of applications that can be extrapolated in a better clinical use.

Even thought, this tool still needs a lot of work and effort to be a more reliable source of STLs for customizable prosthetics hands, we hope that providing the code in an open source platform like github it can be modified to improve and serve his clinical applicable purpose.

## V. CONCLUSIONS

Understanding the affordability of 3D printing technologies and the way of how these can impact the life of patients that need a prosthetic device. It is possible to infer that one of the main limitations to spread the 3D printing of personalized devices is the difficulty of ease customization of them^1, 13, 17^.

Since, many willing medical providers can access the 3D printing devices but they lack on the skills to personalized the prosthetic to each patient in need, and hire specialized personnel to fulfill this task could increase the cost in great manner^1, 6^-^8, 16^.

Here, we propose that a GUI for customization of 3D prosthetic models (trans-radial hand) would be a beneficial contribution to clinical practice in developing countries where a 3D print construct to be efficient must be cheap. Also, a 3D printed prosthetic device to be totally functional must be personalized to everyone. Now, we provide a tool that helps to achieve this goal and allows non-trained medical staff can easily provide 3d customizable prosthetics to all the patients that need it.

